# The hominoid-specific gene DSCR4 is involved in regulation of human leukocyte migration

**DOI:** 10.1101/176503

**Authors:** Morteza Mahmoudi Saber, Marziyeh Karimiavargani, Nilmini Hettiarachchi, Michiaki Hamada, Takanori Uzawa, Yoshihiro Ito, Naruya Saitou

**Author notes:** **Corresponding Author:** Naruya Saitou, Division of Population Genetics, National Institute of Genetics, Yata 1111, Mishima, 411-8540, Japan, /6789. These authors contributed equally to this work (Morteza Mahmoudi Saber, Marziyeh Karimiavargani).

## Abstract

DSCR4 (Down syndrome critical region 4) is an orphan retrotransposon-derived de-novo originated protein coding gene present only in hominoids (humans and great apes). Despite being located on the medically critical genomic region and abundance of evidences indicating its functionality, the role of this gene in human cells was utterly unknown. Due to absence of any prior knowledge regarding the function of DSCR4, for the first time here we used a gene-overexpression approach to discover biological importance and cellular roles of this gene. Our analysis strongly indicates DSCR4 to be mainly involved in regulation of the interconnected biological pathways related to cell migration, coagulation and immune system. We also showed that the predicted biological functions are consistent with tissue-specific expression of DSCR4 in migratory immune system leukocyte cells and neural crest cells that shape facial morphology of human embryo. Immune system and neural crest cells are also shown to be affected in Down syndrome patients who suffer from the same type of DSCR4 misregulation as in our study which further support our findings. Providing evidence for the critical roles of DSCR4 in human cells, our findings establish the basis for further investigations on the roles of DSCR4 in etiology of Down syndrome and unique characteristics of hominoids.

## Introduction

Down syndrome (DS) is a genetic disease with the high incidence of 1 out of 700 live births which turns this disorder into the leading genetic cause of mental retardation and congenital heart disease ^1,2^. The phenotypes characterizing DS are mainly variable in age of onset, frequency and severity and includes immune deficiency, heart disease, dysmorphology of facial characters and underlying skeleton, Hirschsprung’s disease, alterations of brain structure, early onset of Alzheimer pathology and increased risk of leukemia. Mental retardation, characterized by certain behavioral and cognitive deficits is also common feature in DS individuals^3–5^.

DS is caused by the inheritance of an extra copy of the chromosome 21 long arm (q arm). Since the completion of human genome project, it is known that this region span ∼33.5 million base pairs (Mb) of DNA and contain ∼300 genes ^6^. So far mouse transgenic models have been the main tool for the analysis of gene-phenotype correlation in DS ^7,8^. Studies of trisomic mouse models containing an extra copy of mouse genome, orthologous to q arm of chromosome 21, implies upregulation of all trisomic genes by ∼50% across multiple tissues ^7,8^. Although the underlying mechanism accounting for how the relatively small increment in transcription results in any of the commonly observed phenotypes in DS is yet unknown, the correspondence of shortest genomic region shared by DS individuals with the same DS characteristics has led to the hypothesis that a critical chromosomal region called Down Syndrome Critical Region (DSCR) contain a dosage-sensitive gene or set of genes misregulation of which are responsible for emergence of DS features ^9^. The best-defined DSCR spans about ∼5.4 million Mb ranging from a proximal boundary between DS21S17 and D21S55 to distal boundary between MX1 and BCEI ^10^. This region harbor about 33 conserved genes between human and mouse ^11^ and have been associated with multiple DS characteristics such as craniofacial abnormalities, joint hyperlaxity and mental retardation ^2^.

DS Studies using multiple transgenic mouse models in which DSCR orthologous region is overexpressed/ underexpressed, in particular chromosome 16 segmental trisomies such as Ts1Cje, Ts65Dn and Ts1Rhr and segmental monosomies including Ms1Rhr, have so far provided valuable information in constructing the phenotype map of DSCR ^12^. However, investigations on functional annotation of orthologous genes located on DSCR revealed that, first, the mouse orthologous genes are actually dispersed within the genome and do not have the same synteny as human ^13^ and second, not all the genes present in human DSCR region do have orthologs in mouse which proves the inaccuracy of DS mouse models in simulating human Down syndrome condition in mouse.

Down Syndrome Critical Region 4 (DSCR4) is one of the novel de-novo originated protein coding genes on DSCR region which is present only in humans and great apes and lack orthologous counterpart in non-hominine organisms including mice (Fig. 1) ^14^. This gene, also called as DSCRB ^15^ share a bidirectional promoter with DSCR8 gene and is positively and negatively regulated by multiple transcription factors ^16–18^. DSCR4 codes an experimentally verified 118 amino acids long protein ^19^ with strong potentials for forming protein secondary structure ^14^, however, this protein shows no significant sign of homology to any know or putative protein domains ^14,15^. This lack of homology is expected due to the de novo origin of DSCR4 and considering the fact that majority of three coding exons of this gene is derived from retrotransposons ^14^. DSCR4 has been shown be a genetic marker for detection of Down Syndrome ^20^ and hence might act as one of the confounding factors in using mouse models to precisely simulate DS condition. However, no study has so far investigated the functionality and cellular interactions of DSCR4 inside human cells.

**Fig. 1.**
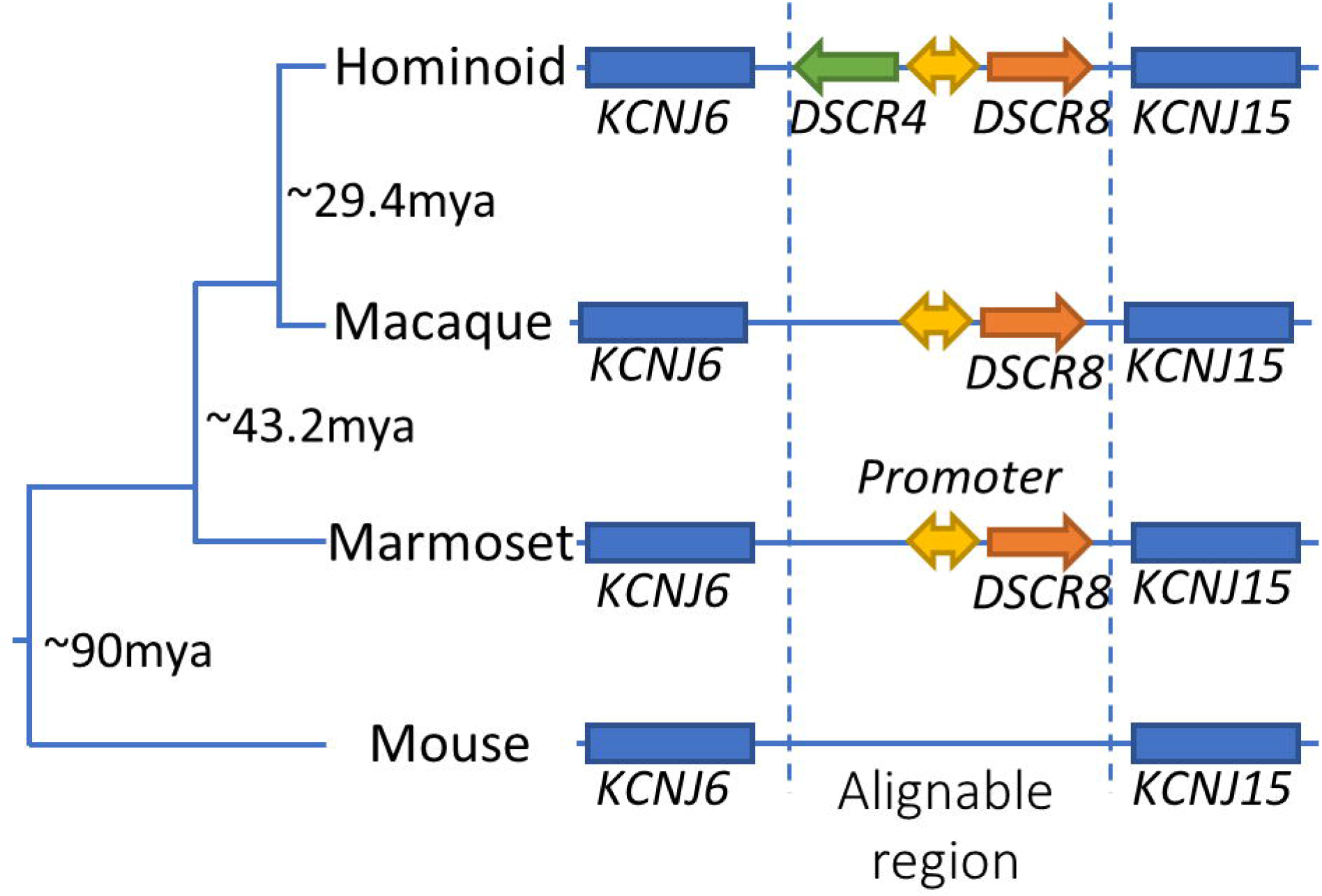
De novo evolution of DSCR4 out of nongenic DNA. Schematic depiction of the evolution of DSCR4 in approximately 29.4 million years ago in common ancestor of hominoids and evolution of DSCR8 and the bidirectional promoter which drives the expression of both genes in approximately 43.2 million years ago in common ancestor of simians.

In this study, to understand the cellular roles and gene regulatory networks in which DSCR4 is involved, we overexpressed the wild type DSCR4 in human non-cancerous cells and measured the consequences on cell transcriptome by differential gene expression analysis. It has been shown that chromosome 21 transcripts are increased proportionally to the gene dosage, i.e. 50% greater than normal in DS cells ^7^. Therefore, to simulate the DS condition, we used overexpression as a means for gene perturbation. Through functional profiling, we investigated the biological processes affected by perturbation of DSCR4 to unravel the role and position of DSCR4 in cellular pathways and its contribution to the etiology of Down syndrome and unique characteristics of hominoids ^21^. Our analysis provides evidences for the functionality of DSCR4 in regulation of cell migration and immune system which are also the biological pathways affected in Down syndrome patients.

## Materials and methods

### Cell culture

Human papillomavirus 16 (HPV-16) E6/E7 transformed cell (HS-27A) which is an immortalized non-cancerous cell type with epithelial morphology were purchased from ATCC^®^ (CRL-2496™) and was cultured in Roswell Park Memorial Institute (RPMI)-1640 medium (Invitrogen) supplemented with 10% fetal bovine serum (FBS) (Invitrogen) according to ATCC^®^ protocols.

### Plasmid DNA vector and control vector construction

PTCN-DSCR4 expression vector (BC096162) which contains full-cDNA sequences of DSCR4 including upstream and downstream untranslated regions were purchased from transOMIC technologies (See supplementary Fig. S1A, Supplementary Material online). No marker was added to DSCR4 cDNA to ensure that the function of the extra copy of DSCR4 is not affected by the marker. Cytomegalovirus (CMV)-derived promoter and enhancer sequences placed before DSCR4 cDNA in the vector ensure proper expression in eukaryotic host cells. For production of control plasmid (PTCN-control), we removed the DSCR4 cDNA sequence along with UTR elements from original plasmid using GeneArt^®^ Seamless Cloning and Assembly kit (See supplementary Fig. S1B, Supplementary Material online). With DSCR4 cDNA being the only difference between PTCN-DSCR4 and PTCN-control, we sought to minimize the confounding elements in our differential gene expression analysis.

### Kill curve assay

The first critical step for generating stably transfected cells is determining the optimal concentration of selection reagent for selecting stable cell colonies. The purpose of kill curve assay is to determine the minimum antibiotic concentration needed to kill all the cells over the course of one week. Since the optimal concentration is cell type dependent, we performed this assay for HS27A cells using cell counting kit-8 (SIGMA). The HS27A cells were treated with G418 (SIGMA) in concentration gradient between 0 (negative control) to 1500 μg/ml. The number of cells in each well were counted after a week and the optimum concentration of G418 for treatment of HS27A cell was determined as 1400 μg/ml.

### DNA transfection and stable cell line selection

For transfection of HS27A cells with PTCN-DSCR4 plasmid, first the optimized concentration of OMNIfect™ transfection reagent was determined using a concentration gradient of OMNIfect and a pcDNA3 vector containing GFP marker. In the next step, using the optimized concentration of OMNIfect (2 μl/ml), the PTCN-DSCR4 and PTCN-control vectors were transfected into HS27A cells. The single cell colonies successfully transfected with DSCR4 were selected by incubation with RPMI-1640 containing 10% FBS and 1400 μg/ml of G418 (Geneticin) over approximately 21 days.

### Acquisition, preprocessing of microarray data and differential expression analysis

Total RNA was extracted from three PTCN-DSCR4 transfected and three PTCN-control transfected HS27A samples along with one normal HS27A sample using PureLink^®^ RNA Mini Kit (Invitrogen) and RNA quality was assessed and confirmed using Agilent 2100 Bioanalyzer (Agilent Technologies). The isolated RNA was then carried through the Agilent preparation protocol and each sample was hybridized to one SurePrint G3 Human 8x60K v3 GeneChip (Agilent Technologies). Raw data were processed and analyzed using the GeneSpring GX software along with RobiNA package ^22^. Quality assessment procedure was conducted by normalized expression value correlation analysis between the three samples groups (See supplementary Fig. S2, Supplementary Material online). The normalized expression values (log base 2) for each chip were calculated using quantile normalization after background correction by RMA method ^23^ which has been proven to measure expression levels reliably. The gene expression matrix was filtered to exclude probe sets which are either not positive or significant, not uniform, not above background or population outliers. After all, saturated probe sets defined as “Class P” by GeneSpring GX software in PTCN-DSCR4 transfected HS27A samples with at least two-fold change in expression comparing with PTCN-control transfected HS27A samples were determined and used for further gene set enrichment analysis.

### Confirmation of DSCR4 gene perturbation using qPCR

Quantitative real-time PCR was conducted by synthesizing cDNA from 1mg DNA-free RNA using random 6-mer primers and PrimeScript First Strand cDNA Synthesis kit (Takara). Quantitative PCR was performed using CFX96 Touch™ Real-Time PCR Detection System. Expression of DSCR4 and GAPDH housekeeping genes were measured using SYBR Green–based gene expression assay (Applied Biosystems). The primers for QPCR analysis were designed to amplify intron-exon boundaries so to avoid amplification of any potential DNA contamination. GAPDH housekeeping gene was used as endogenous control for the normalization. As expected, no significant difference between the cycle threshold (ct) values was observed for GAPDH across all the samples. Melt curve analysis also confirmed strictly specific amplifications in QPCR assays. The comparative Ct approach was employed by CFX96 Touch™ Real-Time PCR machine for quantification of DSCR4 expression levels and confirmed that DSCR4 is overexpressed approximately 9-fold in PTCN-DSCR4 transfected cells comparing with PTCN-control transfected cells (See supplementary Fig. S3, Supplementary Material online)

### Functional profiling

Differentially expressed probe set IDs in PTCN-DSCR4 transfected HS27A samples were mapped to ENTREZ IDs and KEGG IDs and successfully mapped IDs were submitted to ClusterProfiler software ^24^. The Gene Ontology (GO) terms and Gene Regulatory Networks (GRN) significantly overrepresented in the set of differentially expressed genes were identified using ClustreProfiler package. Up- and down-regulated gene sets were analyzed independently using SurePrint G3 Human 8x60K v3 gene sets as reference. GO and GRNs with significant enrichment (p-value < 0.05) were considered in the analysis. Biological processes enrichment analysis was conducted using the data from Reactome curated pathway database ^25^ and pathway enrichment analysis was performed by enrichment test in the set of manually drawn pathway maps provided by Kyoto Encyclopedia of Genes and Genomes (KEGG) ^26^.

## Results

### Functional analysis

There are multiple biological evidences indicating functionality of the de novo originated protein coding gene, DSCR4 gene such as possessing epigenomic marks for active regulatory region within the promoter, potential secondary structure in DSCR4-coded protein, harboring fetal epigenomic marker for detection of Down syndrome and being the binding site for multiple transcription factors ^14^. To verify the functionality of DCSR4 using an evolutionary approach, we also performed Derived Allele Frequency (DAF) analysis. DAF analysis using 1000 genome project data revealed DSCR4 protein coding sequences to have higher rate of polymorphisms with low-frequency derived alleles compared with DSCR4 intronic sequences *(Fig. 2A).* DSCR4 has not been under selection constraint between chimpanzee and human (Dn/Ds ratio = 0.996) ^14^, however, results of DAF analysis prove that DSCR4 CDSs are under purifying selection in human populations.

**Fig. 2.**
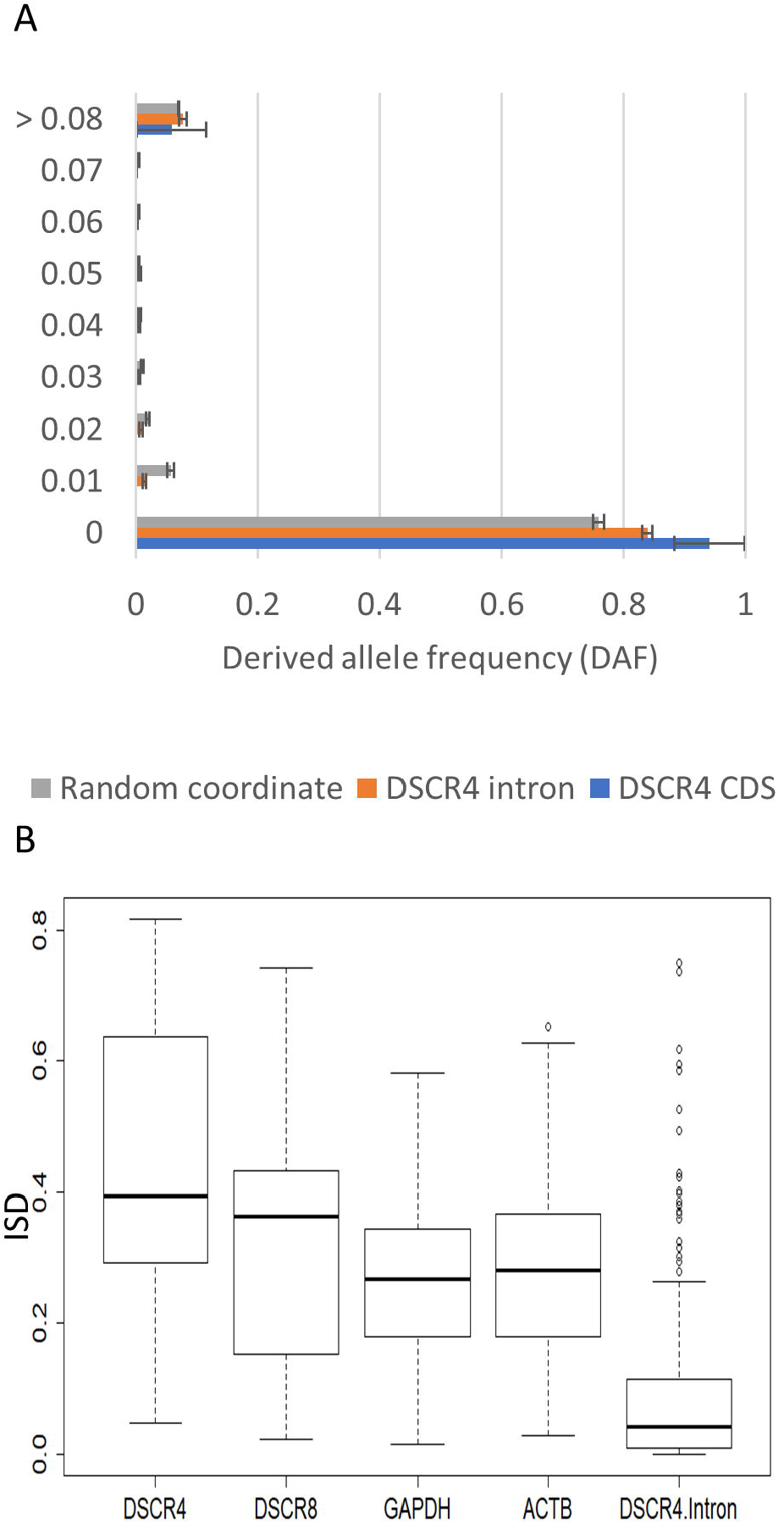
Functional analysis of DSCR4.(A) Derived Allele Frequency (DAF) analysis of DSCR4. DSCR4 CDSs have significantly higher rate of polymorphisms with low-frequency derived alleles compared with DSCR4 intronic sequences indicating the action of purifying selection on the coding sequences of DSCR4. (B) *Intrinsic structural disorder (ISD) characteristics of DSCR4*. DSCR4 protein shows high degree of intrinsic structural disorder comparing to its proximate but older protein coding gene, DSCR8 and conserved old house keeping genes such as GAPDH and ACTB. ISD score of DCSR4 is also significantly higher than that of DSCR4 translated introns.

Intrinsic structural disorder (ISD) is the degree to which a given peptide folds as a stable threedimensional protein i.e. ordered protein versus a rather flexible and unstructured entity i.e. disordered. Natural protein sequences are proved to be more intrinsically disordered than translated random sequences ^27,28^ therefore ISD can be used as a criterion to distinguish functional genes from random ORFs erroneously identified as novel genes. ISD analysis of DSCR4 using IUPred ^29^ indeed revealed that the protein coded by this gene has significantly higher degree of ISD compared with its proximate but older protein coding gene, DSCR8 and conserved housekeeping genes such as GAPDH and ACTB (*Fig. 2*B). Translated DSCR4 intron on the other hand has very low ISD score.

### Differential gene expression analysis

After in silico verification of the functionality of DSCR4, we performed experimental gene perturbation analysis to investigate its functions in vitro. For this purpose, wild-type cDNA and UTR elements of DSCR4 were transferred and overexpressed in immortal human bone marrow cells (HS-27A) cells. DSCR4-transfected human bone marrow cells along with empty-vector (PTCN-control) transfect control HS27 cells were then investigated through transcriptome profiling and differential expression analysis (See materials and methods section). To enhance the reliability of the analysis, the DEGs with unsaturated probe signals which might represent false positives were discarded. We found that a total of 253 probes targeting protein coding genes among 36,427 probes for protein coding genes (<0.5%) are differentially expressed between DSCR4-overexpressing HS27 cells and control samples *(See supplementary Fig. S4, Supplementary Material online)*. Out of 253 differentially expressed genes, 131 represent downregulated and 122 represent upregulated protein coding genes *(Supplementary data, Supplementary Material online)* which were used for further investigations.

### Predicting DSCR4 functions by Gene Ontology analysis

After identification of DSCR4-overexpression-mediated differentially expressed genes, we performed gene ontology analysis to identify the interconnected cellular pathways enriched in DEGs. Functional annotation tools were used to arrange genes in associated categories based on associated gene ontology (GO) terms and participation in biological pathways.

The six GO terms significantly enriched in downregulated DEGs represent interconnected pathways in gene regulatory network of human cells (Fig. 3A and 3B). Four out of the six enriched pathways are directly involved in the regulation of movement, migration and motility of cell or cells compartments. Regulation of cell migration is defined as any process that modulates the frequency, rate or extent of cell migration and regulation of cellular component movement is defined as any process that modulates the frequency, rate or extent of the movement of a cellular component ^30^. Tissue and bone remodeling processes are defined as the reorganization or renovation of existing tissues and bones, respectively. Tissue remodeling can either change the characteristics of a tissue such as in blood vessel remodeling, or result in the dynamic equilibrium of a tissue such as in bone remodeling ^30^. The remodeling processes are critical during development, wound repair and metastatic invasion and are driven by coordinated migration of cells through three-dimensional (3D) extracellular matrix ^31^. The other two pathways in which downregulated DEGs are enriched are also indirectly involved in regulation of cell migration. The pathway, Positive regulation of catenin import into nucleus, is defined as any process that increases the rate, frequency or extent of the directed movement of a catenin protein from the cytoplasm into the nucleus ^30^. The import of β-catenin from cytoplasm to nucleus with assistance of Wnt protein, leads to activation a signaling pathway named Wnt/β- catenin signaling ^32^. It has been shown that Wnt/β-catenin signaling pathway is involved in cell migration of breast cancer cells and metastasis ^32,33^. Negative regulation of JUN kinase is the other enriched pathway in downregulated DEGs and is described as any process that stops, prevents, or reduces the frequency, rate or extent of JUN kinase activity. JUN N-terminal Kinases are a group of kinase enzymes that bind and phosphorylates the c-JUN proteins. C-jun is a proto-oncogene and is the homolog of the viral oncoprotein v-jun ^34^. In breast cancer cells, c-jun is known to play key role in migration and invasion of mammary epithelial cells ^35^. In summary, the enriched pathways and processes within the downregulated DEGs mediated by DSCR4-overexpression consistently indicate the likely role of DSCR4 in regulation of cell migration.

**Fig. 3.**
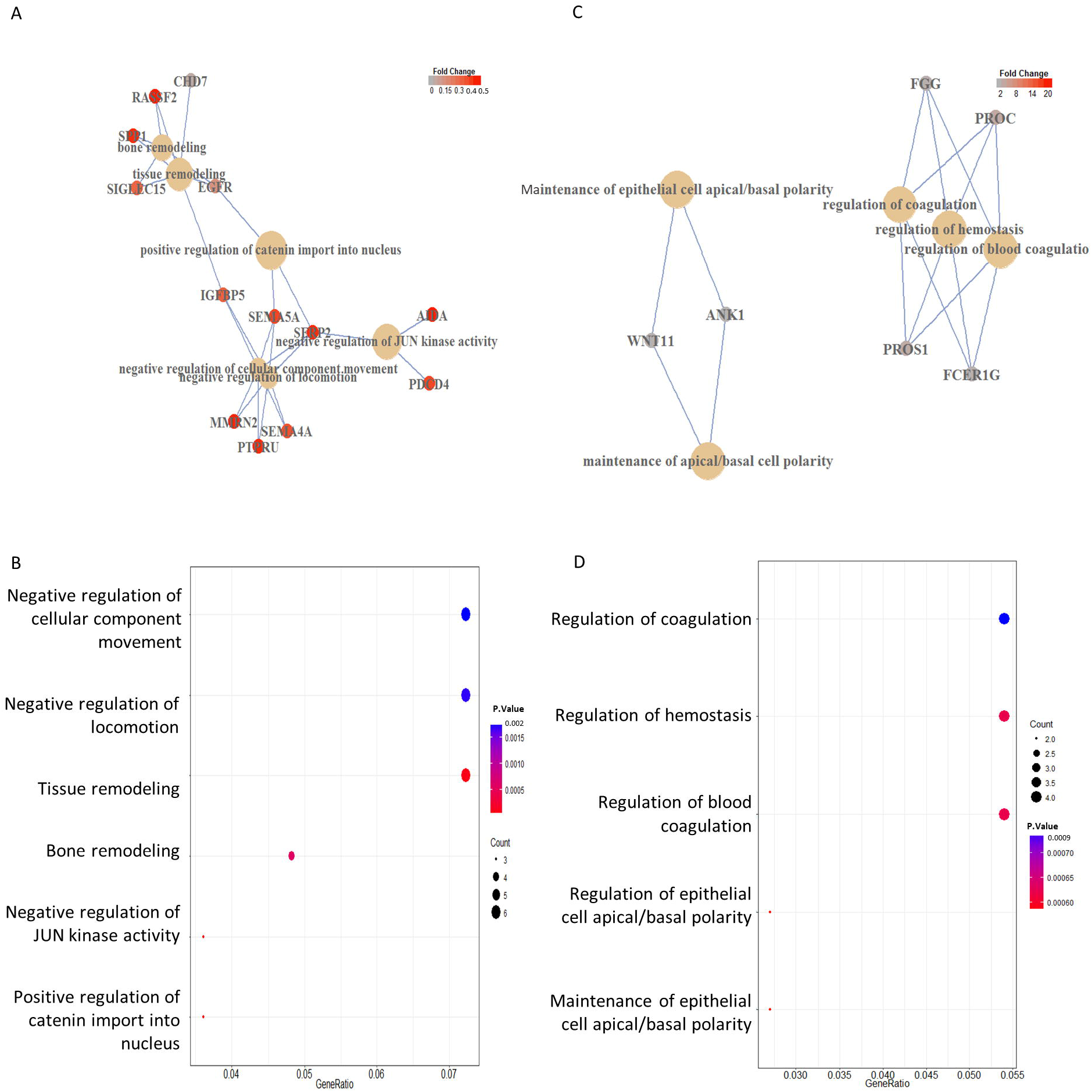
Gene Regulatory Networks (GRN) and Gene Ontology (GO) analysis of DSCR4-overexpression-mediated DEGs. (A) GRN and (B) GO analysis of downregulated DEGs mediated by DSCR4 overexpression, revealed enrichment of 6 interconnected pathways involved in regulation of migration, locomotion and remodeling processes. (C) GRN and (D) GO analysis of upregulated DEGs, on the other hand showed enrichment for three interconnected pathways involved in regulation of blood coagulation and hemostasis along with maintenance of apical/basal cell polarity. (Gene Ratio= number of genes within that list which are annotated to the node / size of the list of genes of interest)

The upregulated genes mediated by DSCR4 overexpression are enriched mainly in regulation of coagulation and hemostasis processes along with maintenance of apical/basal cell polarity (Fig. 3 C and 3D). Regulation of hemostasis and coagulation are respectively defined as the processes leading to stopping of bleeding (loss of body fluid) or the arrest of the circulation to an organ and the processes that modulates the frequency, rate or extent of coagulation. Hemostasis and coagulation processes are critical for the function of blood and innate immune systems ^36,37^. On the other hand, the enriched genes involved in maintenance of apical/basal cell polar, including *wnt11* and *ankyrin1*, were found to be directly involved in regulation of cell migration which is the biological function also enriched in downregulated DEGs. *Ankyrin1* which is a cytoskeleton adaptor protein has been shown to be affected by *p53* and alter cell migration ^38^ and by interacting with silberblick, *Wnt11* is found to be involved in controlling cell migration and morphogenesis in zebra fish ^39^.

### DSCR4-engaged cellular pathways

GO and GRN analysis suggested DSCR4 to be involved mainly in the regulation of cell migration, hemostasis and coagulation. But what are the cellular pathways through which DSCR4 accomplish these functions? To answer this question, we reanalyzed the up- and down-regulated DEGs using the Reactome database data and Kyoto Encyclopedia of Genes and Genomes (KEGG). The reactome databases catalogues a reductionist model which asserts that all the biology can be represented as events located in subcellular compartments ^25^. Consistent with GO and GRN results, these analyses revealed enrichment for a several sub-processes involved in cell migration, coagulation and immune system (See supplementary Fig. S5, Supplementary Material online). Downregulated DEGs are specifically enriched for the function of semaphorines and integrine molecules which are important families of proteins involved in regulation of cell migration (See supplementary Fig. S5A, Supplementary Material online). Strict guidance cues either repulsive, inhibitory or attractive normally precede the process of cell migration and it has been proved that semaphorine receptors govern cell migration mainly by regulating integrin-based cell substrate adhesion and cytoskeleton dynamics ^40^. Semaphorines are also shown to be guiding neural crest cell migration in zebrafish ^41^. Integrins are also a family of membrane molecules important in cell migration and Integrin-based adhesion has served as a model for studying the central role of adhesion in migration ^42^. The upregulated DEGs on the other hand revealed enrichment for the processes of fibrin clot formation and complement cascade which are main columns in coagulation and innate immunity, respectively (See supplementary Fig. S5B, Supplementary Material online). In evolutionary point of view, the two pathways of coagulation and innate immune system have common ancestry and are highly integrated ^43^ which is consistent with our results and support the hypothesis of DSCR4 being simultaneously involved in the function of innate immune system and hemostasis.

Kyoto Encyclopedia of Genes and Genomes (KEGG) pathway database is a collection of manually drawn pathway maps representing existing knowledge on the molecular interaction, reaction and relation networks ^26^. The KEGG analysis further corroborated the results of reactome database by showing that DSCR4-overxpression-mediated DEGs are enriched in coagulation and complement cascade and also it revealed that coagulation and complement cascades are tightly interconnected not only with each other but also with processes of cell migration, adhesion and proliferation (Fig. 4). In summary, these results indicate that all the affected pathways by DSCR4 gene perturbation are closely connected and are mainly involved in the processes of cell migration, immune system and coagulation.

**Fig. 4.**
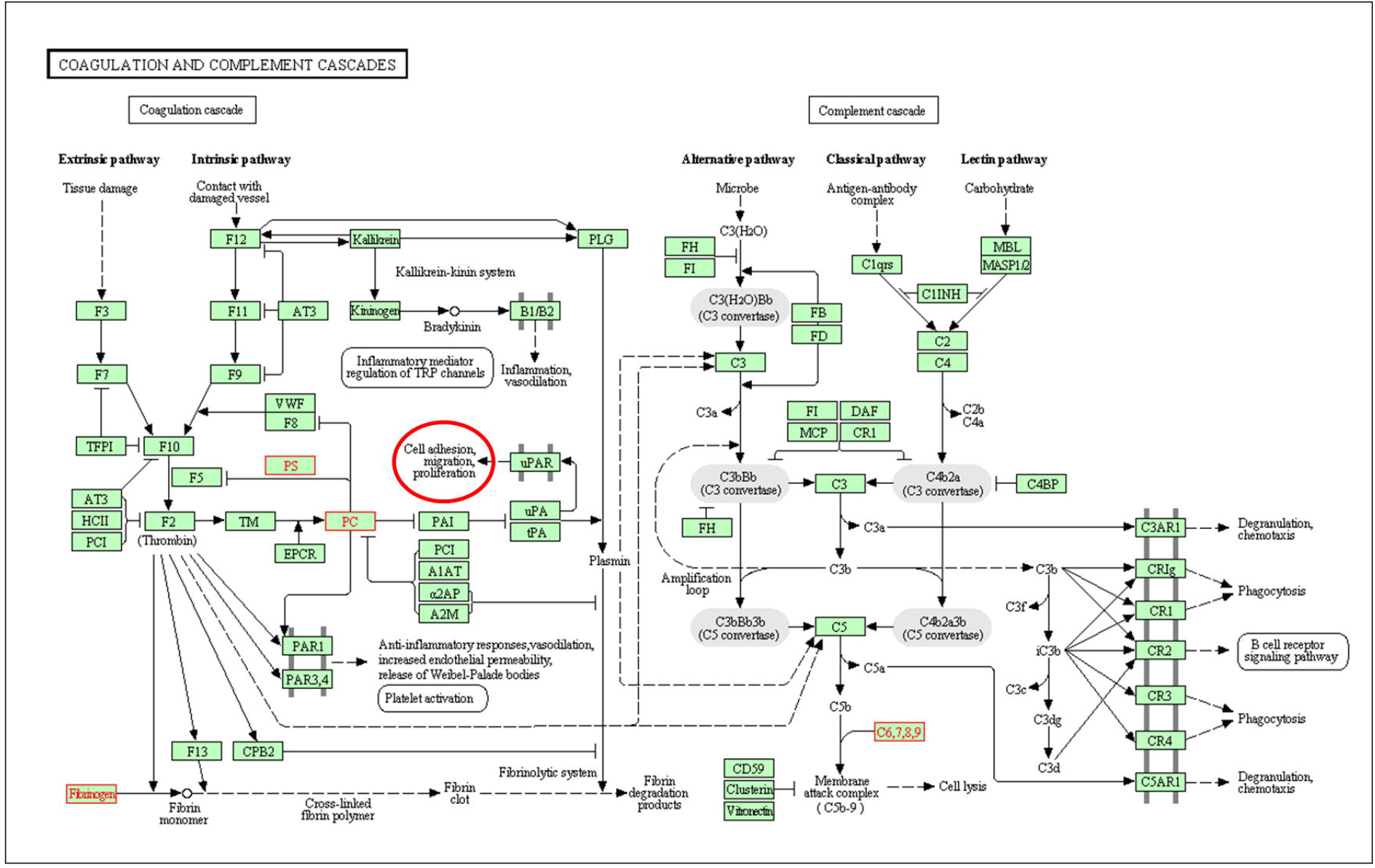
Interconnection of DSCR4-overexpression-mediated perturbed pathways. KEGG analysis of DSCR4-overexpression-mediated DEGs shows the enrichment for tightly interconnected pathways of coagulation cascade and complement cascade (red highlighted) and further confirm the connection of these cascades with cell adhesion, migration and proliferation (red circle).

## Discussion

DSCR4 is an orphan gene present only in Humans and Great apes, evolution of which has occurred in multiple steps over the past 100 million years mainly as a result of the function of retrotransposons (Fig. 1). Young de-novo originated genes have been known with a tendency to be small in size with relatively low expression values and weak conservation signals which also hold for DSCR4 and raise doubts about the functionality of this young de-novo originated experimentally-verified protein coding gene ^18^; recent investigation of intrinsic structural disorder (ISD) however support preadaptation hypothesis of de-novo gene evolution indicating that young genes show extreme levels of gene-like traits and rejects the continuum view of de novo gene birth which suggests a series of intermediate stages, or ‘proto-genes’, between noncoding DNA sequences and a fully functional gene ^44^. Same results were also found in ISD analysis of DSCR4 (Fig. 2B) which is consistent with the results of derived allele frequency analysis (Fig. 2A) and previous findings by Saber et al. ^14^ and further support the functionality of DSCR4.

The results of our gene regulatory network and gene ontology analysis of DSCR4-overexpression-mediated DEGs along with pathway enrichment investigations consistently indicate the likely role of DSCR4 in cell migration, coagulation and immune system. If these predictions are correct, we would expect DSCR4 to be expressed mainly in cells with migratory characteristics. We investigated this hypothesis by quantifying the expression of DSCR4 across all human cells transcriptome data available in Roadmap epigenome project ^45^ and YuGene database ^46^. Only a small portion of highly specialized cells is able to actively migrate within human body which include leukocytes, stem cells and immune system cells ^47^. In line with our predictions, Roadmap data analysis revealed that DSCR4 has significant expression only in K562 cells (Fig. 5). K562 is human immortalized myelogenous leukemia cell with erythroleukemia type. Consistently, analysis of YuGene database also revealed that monocytes which are a type of white blood cells or leukocytes are the cells with highest expression level of DSCR4 (See supplementary Fig. S6, Supplementary Material online). After, monocytes, T-cells and Dendritic cells (DSs) are the two other immune system cells notably expressing DSCR4 with proven migratory behavior. Three types of stem cells and their derivatives including embryonic stem cells (ESCs), ESC-dervied neuron and induced pluripotent stem cells (IPScs) with migratory features are also among the top cells significantly expressing DSCR4. Blastocysts, the structure formed early in the development of all mammals is yet another type of cell with migratory characteristics significantly expressing DSCR4. Neural crest cells (NCCs) are a group of embryonic cell population emerging early in the development of vertebrates and most relevant to the unique 48 human facial traits. These cells arise from the dorsal part of the neural tube ectoderm and migrate into the branchial arches and what will later form the embryonic face structure, consequently establishing the central plan of facial morphology ^49^. Analyses of YuGene database (See supplementary Fig. S6, Supplementary Material online) along with the NCC transcriptome analysis in human and chimpanzee by Prescott et al. ^48^ (Fig. 5), indeed, revealed the significant expression of DSCR4 in critical and migratory NCC cells as expected. In summary, the expression profile of DSCR4 across human cells and tissues support the predicated functions of this gene by our gene perturbation analysis and indicate the likely role of DSCR4 in regulation of cell migration and immune system.

**Fig. 5.**
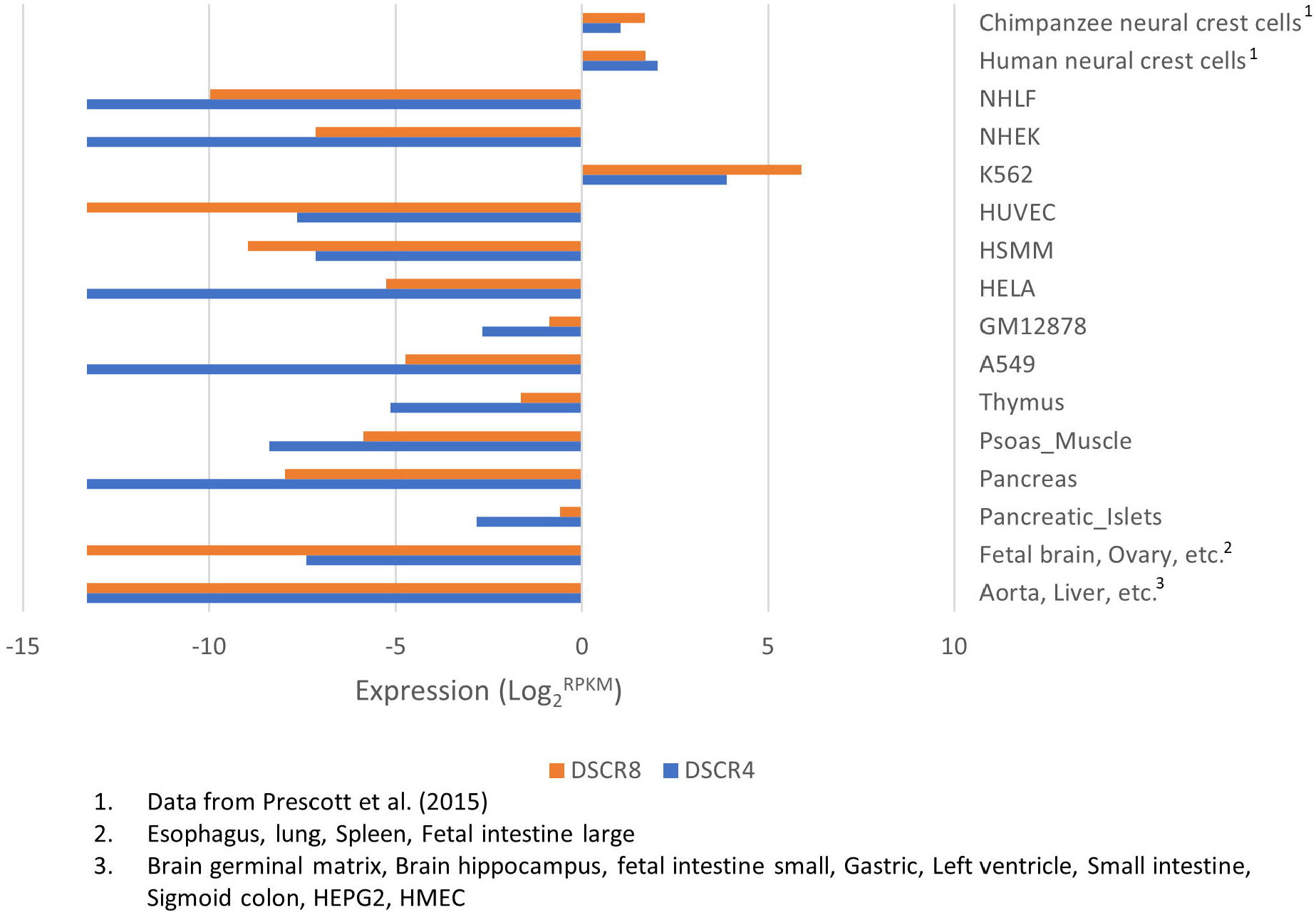
Expression profile of DSCR4 across human cell lines and tissues. According to Roadmap epigenome project data, DSCR4 and DSCR8, which share a bidirectional promoter, do have significant expression only in K562 cells which is a type of leukemia cells. Analysis of transcriptome data provided by Prescott et al. (2015) show that DSCR4 and DSCR8 do also have significant expression in neural crest cells of humans and chimpanzees which are critical migratory cells involved in formation of facial morphology of embryo.

On a deeper level, if DSCR4 is involved in regulation of cell migration, coagulation and immune system, it is expected to observe related symptoms in human disorders with DSCR4 overexpression. Down syndrome is human inherited disorder caused by an extra copy of whole or part of chromosome 21, specially at DSCR region where DSCR4 is located, leading to an average 50% overexpression of genes located on this region. Acute leukemia has been associated with Down Syndrome 10-20 fold higher than general population and this disorder is known as one the most leukemia-predisposing syndrome ^50,51^. Additionally, DSCR4 has been shown to be under copy number amplification in leukemia cells (See Supplementary Fig. S7, Supplementary Material online) ^52^. Leukocytes are top human cells expressing DSCR4 to significant levels (Fig. 5 and Supplementary Fig. S6, Supplementary Material online) with active migratory characteristics and 53 prominent role in coagulation process which are the biological functions enriched in DSCR4-overexpression-mediated DEGs. Another main characteristic of Down Syndrome patients is immune deficiency, especially at innate immune system ^54^. This symptom of Down syndrome patients is consistent with our findings as innate immune system is enriched with genes affected by DSCR4 overexpression (Fig. 4 and Supplementary Fig. S5, Supplementary Material online). In addition, multiple immune system cells including monocytes (leukocytes), T-cells and dendritic (DS) cells were shown to be notably expressing DSCR4 gene (See Supplementary Fig. S6, Supplementary Material online) which indicate active role of DSCR4 in immune system cells. Dysmorphology of facial characteristics is yet another main feature of Down Syndrome patients which can be explained, at least in part, by our findings that DSCR4 is involved in regulation of cell migration and that neural crest cells (NCC) do have notable expression of DSCR4 (Fig. 5 and Supplementary Fig. S6, Supplementary Material online). Cell migration is essential for the function of NCCs which form the facial characteristics of embryo, hence, imbalanced DSCR4 expression can affect the migratory behavior of NCCs which in turn could affect facial morphology of Down syndrome patients.

DSCR4 is an orphan retrotransposon-derived gene existing only in humans and great apes which although locating a medically critical region, its function was yet unexplored. Because there was no previous study on the functions of this gene, we were obliged to use a brute-force approach for identification of its functions that is a statistically challenging objective considering the moderate cellular effects of short and young genes. The results of our gene perturbation and differential expression investigations however are consistent with transcription profile of human cells and human diseases with DSCR4 gene perturbation indicating the likely role of DSCR4 in interconnected pathways of cell migration, immune system and coagulation. Our findings further indicate the importance of retrotransponses in genetic innovation within hominoid genome and for the first time, here we provide solid predications on the functions DSCR4 which can be subject of future direct investigations to unravel the details of the functionality and roles of this orphan and yet unclassified gene in etiology of Down syndrome and other disorders such as cancer metastasis.

## Acknowledgments

This work was supported by a research support grant from Sasakawa Company and foreign student fellowship from Ministry of Education, Culture, Sports, Science and Technology (MEXT) of Japan to M. M. S. and by a grant-in-aid for scientific research from MEXT of Japan to N.S. Some of the analyses were performed on National Institute of Genetics of Japan supercomputer.

